# Atomic structures determined from digitally defined nanocrystalline regions

**DOI:** 10.1101/820274

**Authors:** Marcus Gallagher-Jones, Karen C. Bustillo, Colin Ophus, Logan S. Richards, Jim Ciston, Sangho Lee, Andrew M. Minor, Jose A. Rodriguez

## Abstract

Nanocrystallography has transformed our ability to interrogate the atomic structures of proteins, peptides, organic molecules and materials. By probing atomic level details in ordered sub-10 nm regions of nanocrystals, approaches in scanning nanobeam electron diffraction extend the reach of nanocrystallography and mitigate the need for diffraction from large portions of one or more crystals. We now apply scanning nanobeam electron diffraction to determine atomic structures from digitally defined regions of beam-sensitive peptide nanocrystals. Using a direct electron detector, we record thousands of sparse diffraction patterns over multiple crystal orientations. We assign each pattern to a specific location on a single nanocrystal with axial, lateral and angular coordinates. This approach yields a collection of patterns that represent a tilt series across an angular wedge of reciprocal space: a scanning nanobeam diffraction tomogram. From this diffraction tomogram, we can digitally extract intensities from any desired region of a scan in real or diffraction space, exclusive of all other scanned points. Intensities from multiple regions of a crystal or from multiple crystals can be merged to increase data completeness and mitigate missing wedges. Merged intensities from digitally defined regions of two crystals of a segment from the OsPYL/RCAR5 protein produce fragment-based *ab-initio* solutions that can be refined to atomic resolution, analogous to structures determined by selected area electron diffraction. In allowing atomic structures to now be determined from digitally outlined regions of a nanocrystal, scanning nanobeam diffraction tomography breaks new ground in nanocrystallography.

## Introduction

A prominent bottleneck to the determination of atomic molecular structures is their formation of well-ordered single crystals of a suitable size. As a crystal grows, so too does its likelihood of being disordered^1^. Lattice disorder can result in the loss of diffracting power, challenges in data reduction and ultimately increases the difficulty of structure determination^2^. Microfocus X-ray beams overcome some of these challenges, reducing the lower size limits of crystals from 100s of microns to below 10 microns^3^. Serial crystallography at both synchrotron^4^ and X-ray free electron laser sources^5^ further reduces crystal size limits to the sub-micron scale, at the cost of requiring large numbers of crystals. Electron diffraction has undergone a recent renaissance in the form of microcrystal electron diffraction (MicroED or cRED)^6–8^, which allow the structures of protein^6^ or organic small molecules^9,10^ to be determined from 3D nanocrystals.

Each of these advances has revealed novel structures: G-protein coupled receptors first determined at microfocus beamlines^11^, cell grown crystals were interrogated by XFEL beams^12^, whilst MicroED has revealed high-resolution structures of the toxic cores of many amyloidogenic proteins^13,14^. MicroED has also proven to be a powerful method for the interrogation of small molecule structures, revealing atomic structures from seemingly amorphous powders^9^. Electron nanobeams^15^ approximately 2 - 150 nm in size can facilitate diffraction from challenging beam sensitive materials such as zeolites^8^, polymers^16,17^, organic small molecules^18^ and proteins^19^, as well as more radiation hardy inorganic materials^20^.

Capitalizing on innovations in electron nanodiffraction^21,22^, we demonstrate the collection of high-resolution tomographic diffraction tilt series from single crystals using electron nanobeams with a full width at half maximum of ∼12 nm. Scanning nanobeam electron diffraction tomography (nanoEDT) data is collected by coupling four dimensional scanning transmission electron microscopy (4DSTEM) strategies^23,24^ with sample tilting along one or more axes. Meaningful diffraction signal is measured using a hybrid counting strategy implemented on sparse data collected on direct electron detectors. Data is reduced from digitally selected areas of a scan and used in Fourier synthesis to determine the structure of an amyloid-forming segment of the OsPYL/RCAR5 protein from *Oryza sativa*, a rice abscisic acid receptor. The determination of this peptide structure by nanoEDT severs our need for use of a pre-defined diffraction aperture during data collection and opens a new realm of possibilities for structure determination from arbitrarily defined nanocrystalline regions.

## Results

To assess whether meaningful diffraction could be collected from nanometer size regions of a crystal by nanoEDT, we scanned a focused electron beam of 12 nm in diameter (Sup. Fig. 1) through crystals of the OsPYL/RCAR5 peptide. The crystals were needle shaped (Sup. Fig. 2-3) and approximately 360 nm thick, 500 nm wide and several micrometers in length (Fig. 1, Sup. Fig. 4). In nanoEDT, a tilt series was collected as consecutive scans at specified angles, typically separated by one to two-degree increments (Sup. Fig. 2-3). Each scan grid had a spacing of 40 nm covering a total area of 1 by 4 µm (Fig. 1). The spacing in our scans ensured minimal probe overlap between adjacent illuminated areas in each scan, thus limiting the total dose imparted across the crystal. The diffraction patterns from a single scan at a single crystal orientation were then computationally combined to produce a single diffraction pattern that represented the sum of all electron counts across a defined region of the scan (Fig. 1).

**Fig. 1.**
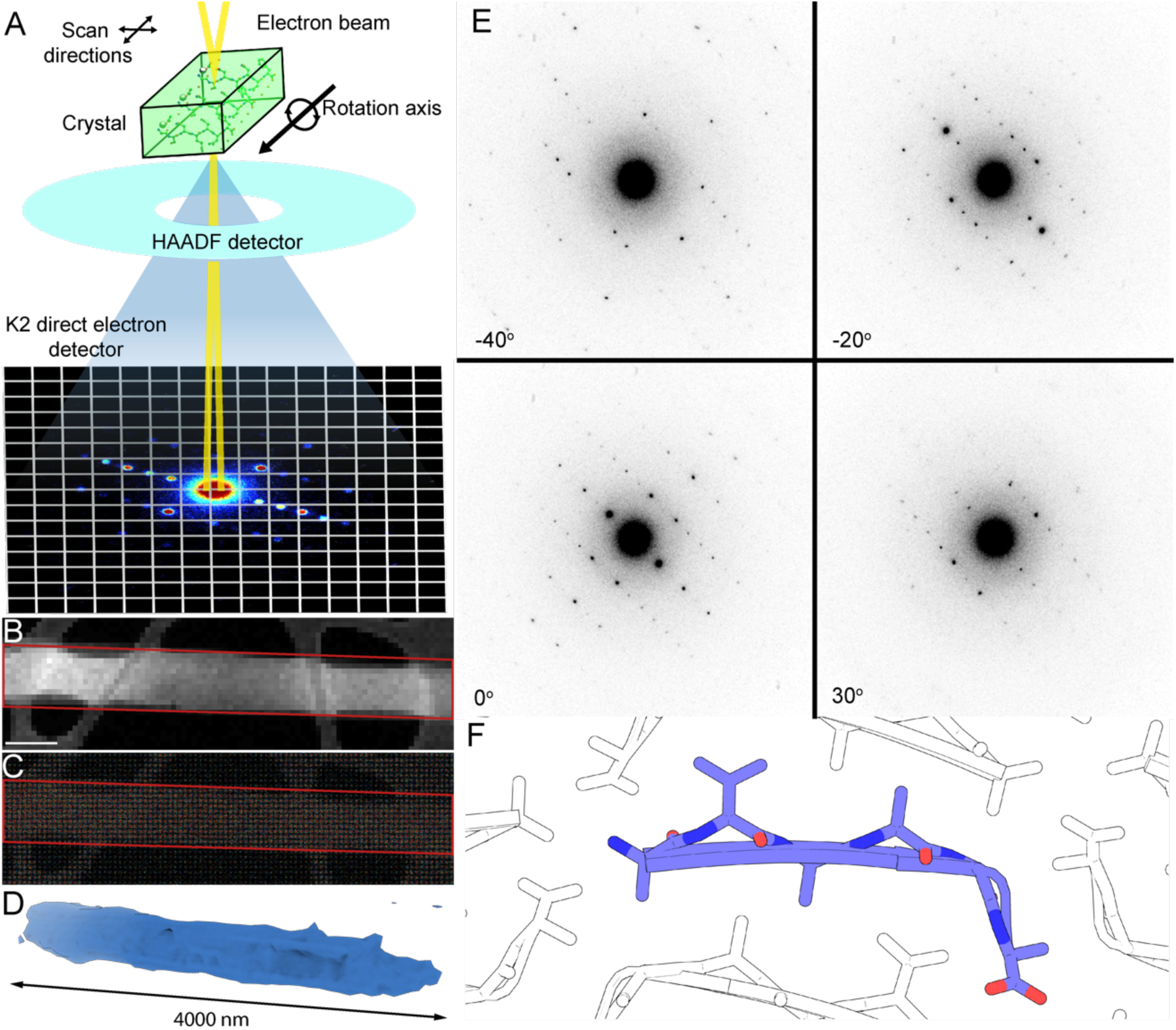
Overview of scanning nanobeam electron diffraction tomography (nanoEDT) experiment. (A) Schematic of experimental geometry for collecting nanobeam electron diffraction data with key components highlighted. (B) ADF image of a crystal of segment ^24^AVAAGA^29^ from the OsPYL/RCAR5 protein interrogated by an electron beam. Scale bar represents 400 nm. (C) Composite image of all diffraction patterns collected simultaneously with the ADF image in (B). Red outline indicates region of the image used to compute diffraction patterns. (D) Tomographic reconstruction of the crystal in (B). (E) Examples of diffraction images taken at discrete orientations during electron diffraction tomography. (F) Atomic structure of OsPYL/RCAR5 peptide ^24^AVAAGA^29^ solved by nanoEDT.

By exploiting nanoEDT’s ability to construct a real space image from the diffraction data, we could digitally select diffraction from a specified region of a crystal or field of view. From the annular dark-field (ADF) image acquired simultaneously with the diffraction patterns or from a reconstructed virtual darkfield image we were able to identify and segment regions of interest in each scan. Diffraction signal was then selected only from these regions to produce a final set of diffraction patterns. The regions encompassed the distinguishable bounds of the crystal (Fig. 1., Sup. Fig. 2 and 3); their dimensions match those obtained from three-dimensional tomographic reconstructions of target crystals based on estimates of their thickness (Sup. Fig. 4, 5). By rotating the sample stage one degree between scans, we computed 81 summed diffraction patterns spanning an angular range of ±40^0^ (Fig. 1). The nominal exposure over the full rotation series is ∼81 e^-^/Å^2^, within the range of a typical tomography experiment^25^.

We indexed and integrated nanoEDT data from regions of interest in two different crystals of the OsPYL/RCAR5 peptide, and then assembled tilt series from each crystal into a 3D reciprocal lattice using conventional crystallography software. The outermost reflections observable in each tilt series corresponded to ∼1.1 Å resolution (Sup. Fig. 5) and an overall completeness of 70% at 1.35 Å (table 1). While the diffracted signal at high-resolution was not sufficiently complete for direct methods, *ab-initio* fragment-based phasing using the program Arcimboldo was successful in generating initial phases from a library of probes consisting of poly-glycine tetramers (Fig. 2). A single 4-residue β-strand was placed by the Arcimboldo suite of programs^26,27^ with a log likelihood gain (LLG) of 35.9 and an initial correlation coefficient (CC) of 55.49. This was subsequently built and refined against electron atomic scattering factors in Phenix^28^ (Fig. 2) to produce a class 4 amyloid zipper^29^ with 6 residues per strand. The refined structure yielded final crystallographic *R*-factors of *R*_work_/*R*_free_: 0.253/0.260. This was consistent with structures solved by microED with a comparable level of electron exposure (Table 1). The overall structure had *B*-factors that were sufficiently low (1 – 5 Å^2^) to detect hydrogen atoms for many of the residues at the core of the zipper (Fig. 2). Residues at the C-terminus showed considerably higher *B*-factors than the rest of the structure resulting in less well-defined density in this region (Fig. 2). We initially attempted to model hydrogen atoms at this position, however in doing so, the *R*-factors slightly increased, and the density around the C*α* carbon significantly depreciated leading us to exclude these hydrogen atoms in the carboxy terminus of the final model.

**Table 1.**
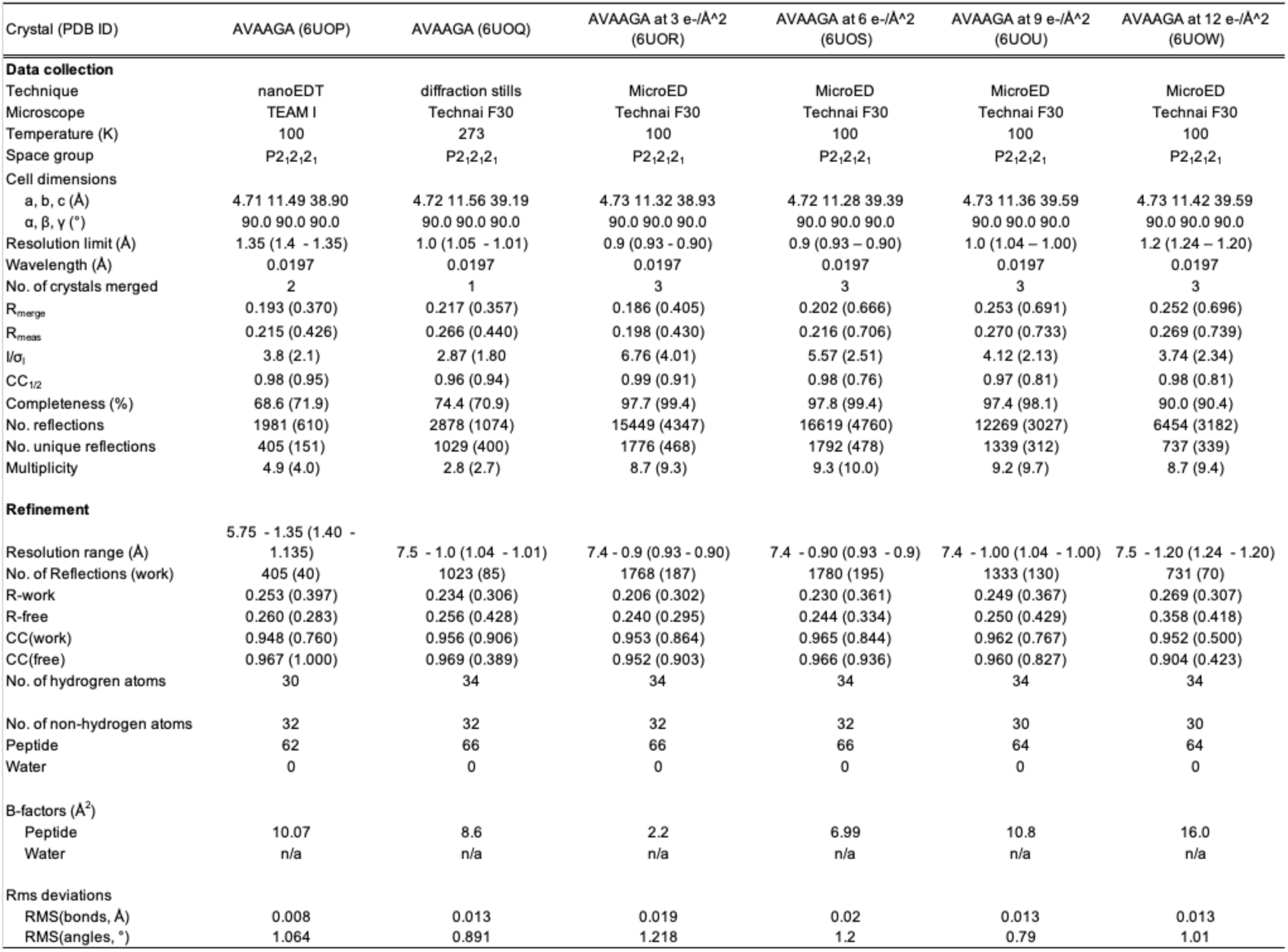
Data collection and refinement statistics.

**Fig. 2.**
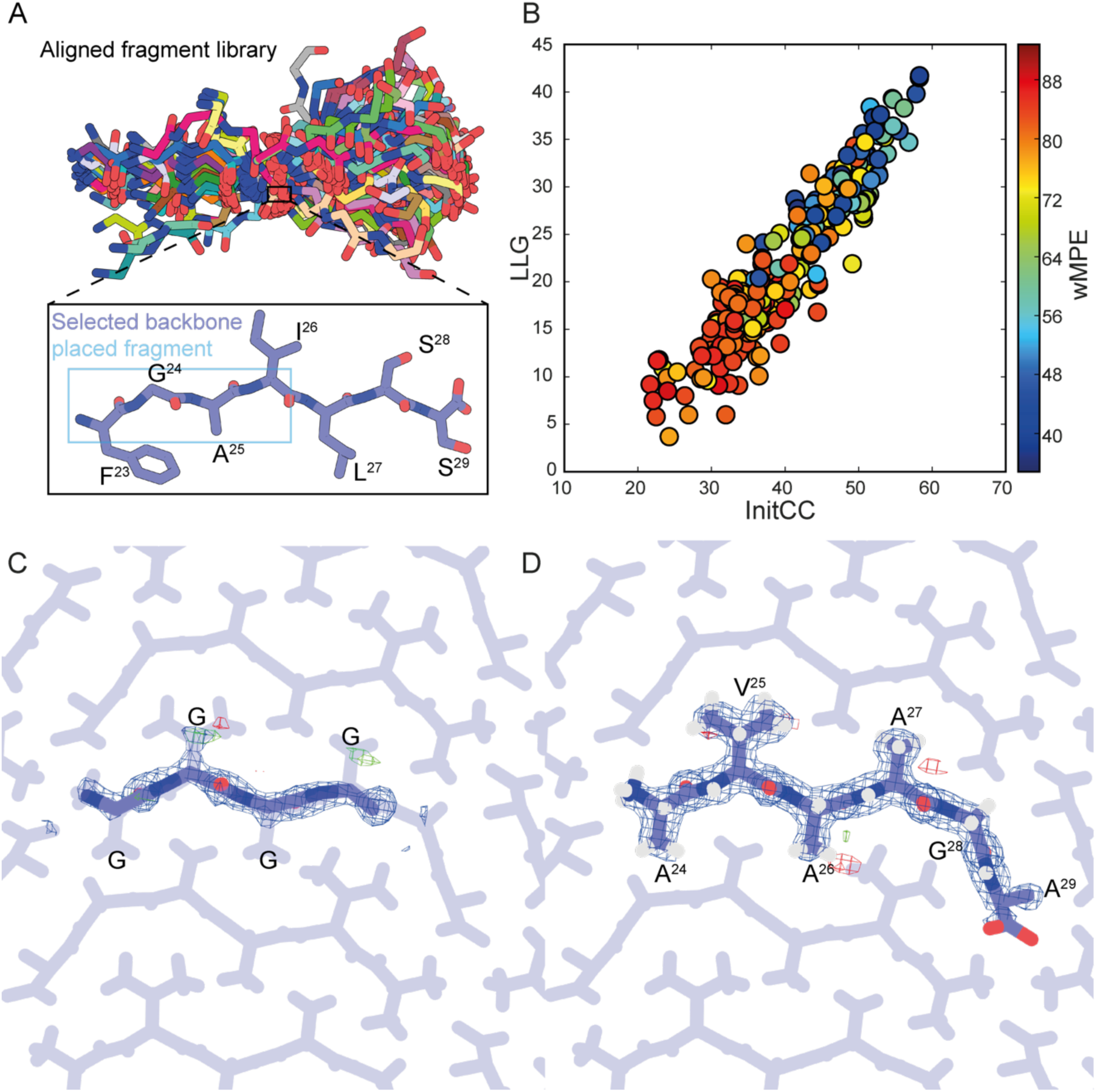
Fragment-based phasing of nanoEDT data. (A) Amyloid peptide fragment library used as input for Arcimboldo. The final fragment placed and the structure it is derived from^32^are highlighted by the blue and black boxes respectively. (B) LLG *vs.* Initial CC for all fragments used by Arcimboldo to find initial phasing solution. The color bar represents the mean phase error of a given fragment compared to the final solution. (C) Initial fragment placed by Arcimboldo (blue) overlaid on the final solution (purple). (D) Final refined structure of OsPYL/RCAR5 peptide ^24^AVAAGA^29^. Blue mesh represents the 2Fo-Fc map (contoured at 1 *σ*) and green/red mesh represents the Fo-Fc map (contoured at ±3 *σ*).

We assessed the accuracy of intensities measured by nanoEDT by comparing them to intensities measured by selected area electron diffraction approaches: either continuous rotation MicroED or fixed-angle diffraction. We determined structures of the OsPYL/RCAR5 peptide from diffraction collected by continuously rotating crystals in an electron beam (MicroED), and by capturing diffraction at fixed angles in discrete 1° increments from crystals whilst exposing them to a 300 kV electron beam. All experiments were performed on crystals of the OsPYL/RCAR5 peptide from the same batch condition and prepared in the same way; in all cases the angular sampling was ±45 degrees. Structures were determined by direct methods from 2 different datasets: merged MicroED data from 3 crystals and fixed-angle diffraction recorded from a single crystal. Comparison of the structures from all 3 datasets showed a high degree of similarity with an overall all atom RMSD of 0.145 ±0.03 Å with the greatest deviation occurring at the C-terminus (Fig. 3). The overall statistics of the refinements are summarized in Table 1. The best quality data was obtained by merging MicroED diffraction data from several crystals, as reflected in the final refined *R*-factors. Interestingly we note that the *R*-factors observed from both nanoEDT data and fixed-angle diffraction are similar to those obtained from conventional MicroED data despite potential issues with partiality (Table 1).

**Fig. 3.**
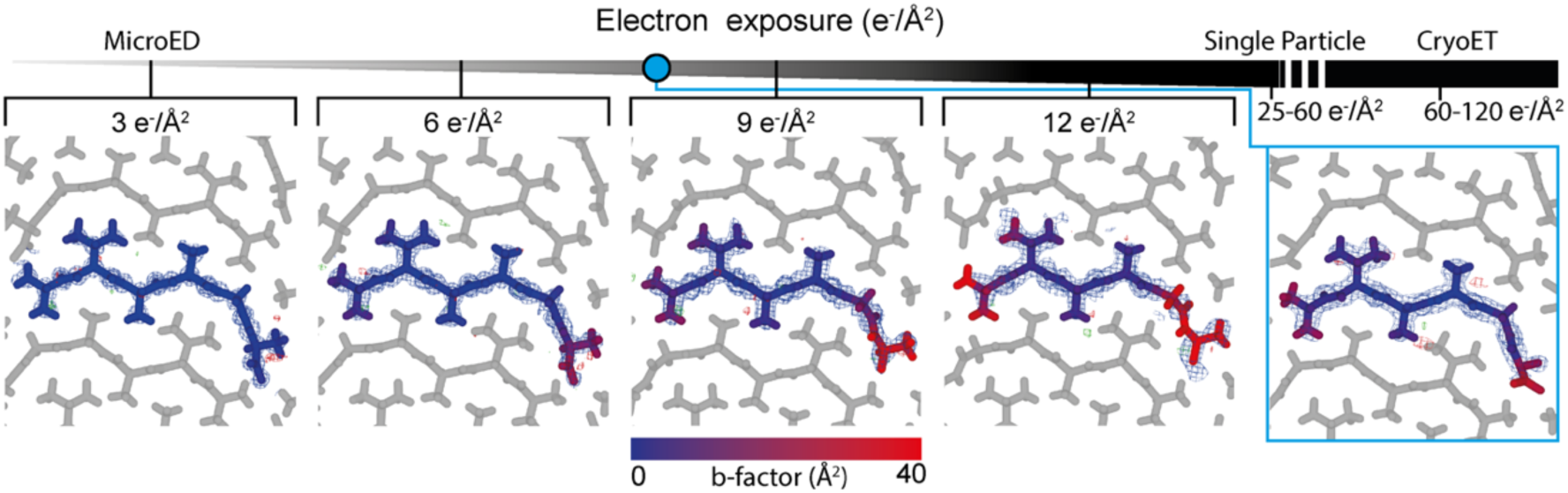
Estimation of electron exposure in nanoEDT. The top gradient represents increasing exposure to the incident electron beam. Several cryo-EM methods are highlighted with typical values of exposure. The blue dot indicates the apparent exposure of the nanoEDT structure based on the comparison to observed b-factors in structures solved by MicroED at a known electron exposure. Blue mesh represents the 2Fo-Fc map (contoured at 1 *σ*).

To further explore differences between the various ED datasets, we performed pairwise comparisons of the magnitudes from each dataset after scaling their intensities. We performed linear regression on these comparisons to visualize and quantify the correlation between datasets (Fig. 3). Overall, the fixed-angle diffraction data had the poorest correlation to all other datasets, with the highest correlation being between the data taken by conventional MicroED and nanoEDT (Fig. 4). Visual inspection of the distribution of Bragg peak intensities across the three principle zone axes in all datasets supported this high degree of similarity (Sup. Fig. 5). However, comparisons along the 0KL and H0L zone axes were limited by the narrow wedge of data collected by nanoEDT, exacerbated by the orientation bias of OsPYL/RCAR5 peptide crystals on the grid in these experiments.

**Fig. 4.**
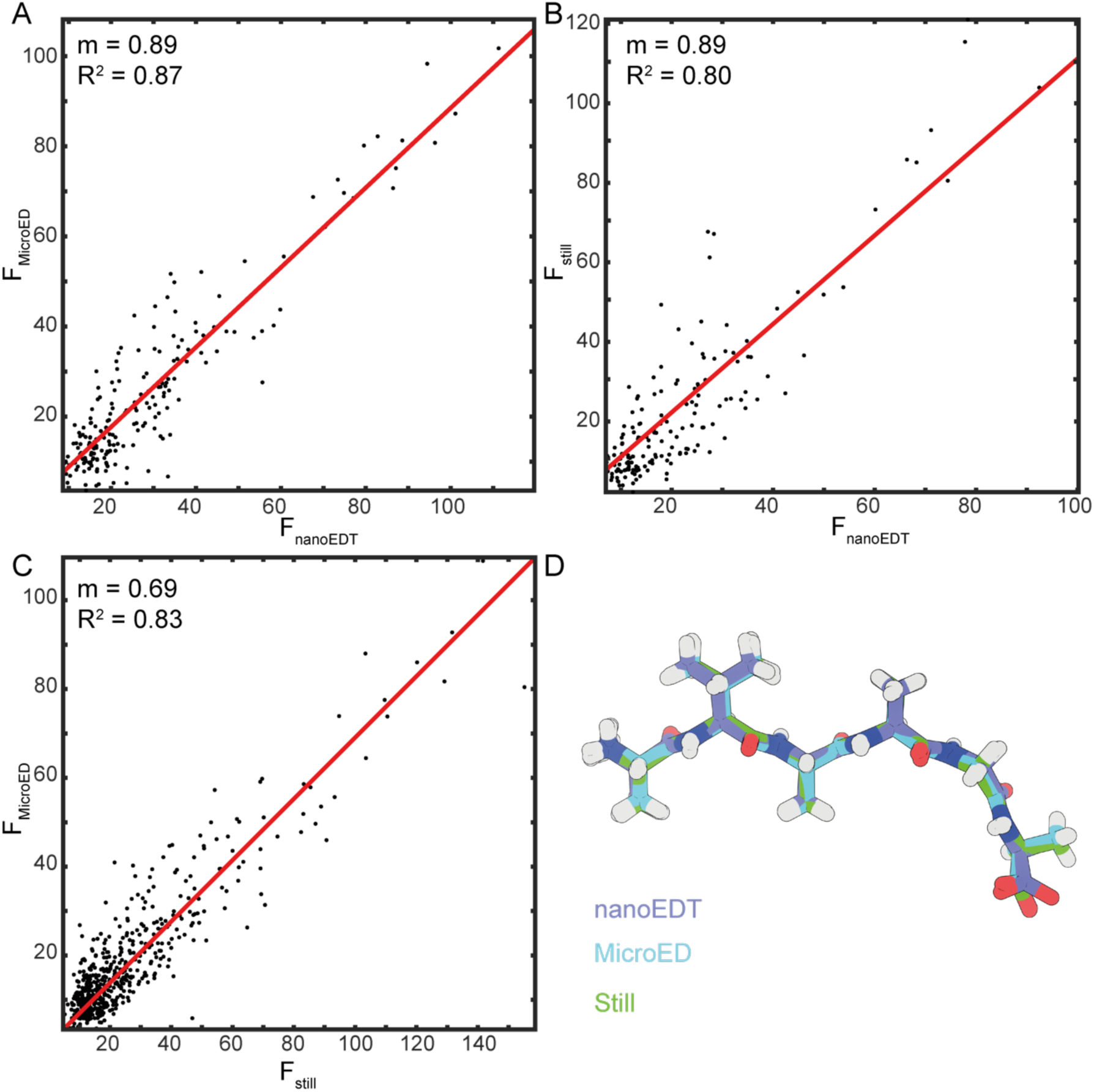
Pairwise comparison of Fourier magnitudes of OsPYL/RCAR5 peptide ^24^AVAAGA^29^ crystals recorded by different methods. (A) Linear regression fit to the pairwise comparison of Fourier magnitudes collected using MicroED and nanoEDT. (B) Linear regression fit to the pairwise comparison of Fourier magnitudes collected using fixed-angle diffraction and nanoEDT. (C) Linear regression fit to the pairwise comparison of Fourier magnitudes collected using MicroED and fixed-angle selected area diffraction. (D) Alignment of the structures determined by each of the three methods. The all atom RMSD is < 0.15 Å.

Although the exposure per illuminated region in nanoEDT is considerably higher than in microED, the observed impacts of its higher exposure on the final structure determined by nanoEDT are more consistent with a conventional microED exposure^30^. To evaluate the impact of electron exposure during nanoEDT data collection, we compared our nanoEDT structure of the OsPYL/RCAR5 peptide to those determined by MicroED under various exposures at cryogenic conditions to a 300 kV electron beam. To observe the effect of increasing electron exposure on OsPYL/RCAR5 peptide crystals we collected 4 consecutive datasets from 3 different crystals with a total estimated exposure of 3 e^-^/Å^2^/dataset. Merging the data from 3 different crystals allowed us to determine MicroED structures of OsPYL/RCAR5 peptide with a collective exposure of 3, 6, 9 or 12 e^-^/Å^2^ (Fig. 4). We observed that as exposure increases, there is a proportionate increase in the *B*-factors of the atoms at the carboxy-terminus of the OsPYL/RCAR5 peptide structure. This was coupled with an overall loss of resolvable density in this region (Fig. 4). By the time the crystals had been exposed to 9 e^-^/Å^2^, the OsPYL/RCAR5 peptide structure showed no visible density for C-terminal oxygens. Because the OsPYL/RCAR5 peptide structure determined by nanoEDT shows a *B*-factor profile in between the 6 and 9 e^-^/Å^2^ exposure structures in the MicroED dose series, we believe the effective dose experienced by the crystals in the nanoEDT is consistent with an effective exposure of 6 to 9 e^-^/Å^2^ (Fig. 4).

The ability to digitally define regions of interest using nanoEDT extends to polycrystalline samples and clustered crystals, from which coincident lattices can be separated yielding high-resolution single crystal diffraction (Fig. 5). This is achieved by integrating diffracted signal from separate regions within adjacent crystallites, allowing the identification of each lattice within a multi-lattice pattern (Fig. 5). This approach relies on spatial separation of crystallite regions in ADF images or simulated dark field images of a grid region, and thus avoids the need for lattice deconvolution or multi-lattice indexing^31^.

**Fig. 5.**
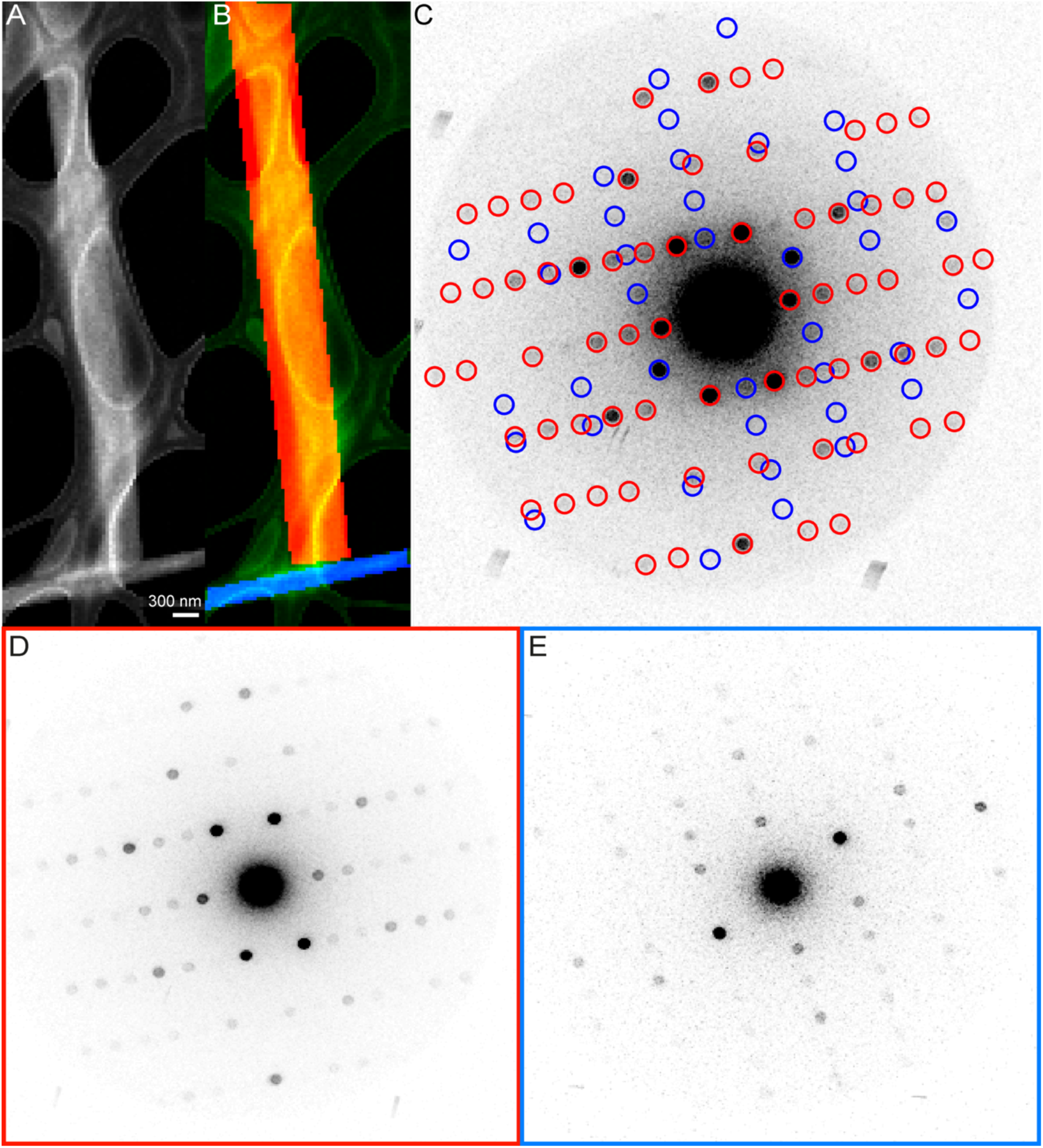
Digital separation and extraction of multiple lattices from separate crystals in a single field of view. (A) ADF image of two OsPYL/RCAR5 peptide crystals. (B) Segmentation of the 2 crystals from (A). (C) 4DSTEM pattern calculated from the entire field of view in (A). Bragg reflections arising from the masked regions in (B) are highlighted by circles of their respective color. (D) 4DSTEM pattern calculated from only diffraction patterns captured from the red region of (B). (E) 4DSTEM pattern calculated from only diffraction patterns captured from the blue region in (B).

## Discussion

In a first demonstration of the powerful application of scanning nanobeam electron diffraction, using a tomographic diffraction approach (nanoEDT), we determine the atomic structure of an amyloid-forming OsPYL/RCAR5-derived peptide phased by fragment-based methods. We demonstrate the capture of meaningful diffraction from regions of a peptide crystal as small as 40 nm and combine this data digitally post-experiment. Subsequent data reduction allows for structural determination and refinement from user selected areas of single or clustered nanocrystals. Structures determined by nanoEDT are accurate, comparing favorably to structures of the same sample determined by selected area diffraction methods, and have refinement statistics comparable to those from other methods (Table 1). However, nanoEDT allows atomic detail to be extracted from a digitally defined nanoscale volume.

The general applicability of nanoEDT to various nanocrystalline substrates is limited only by their diffracting quality at the 10-40 nm scale, which matches the beam sizes and grid samplings demonstrated in our experiments. Our current efforts correspond to observations from a single peptide but the methods implemented could benefit a broad variety of nanocrystalline samples with an equivalent or greater tolerance to electron exposure. We note that the estimated electron exposure (∼81 e^-^/Å^2^) is far greater than the impact observed on the structure determined by nanoEDT, which corresponded best with MicroED structures irradiated 10-fold less. We rationalize this by noting that since scan points were 40 nm apart on a regular grid, the crystalline area mapped in a single scan step (1600 nm^2^) is approximately 14 times larger than the area directly illuminated by the electron beam (113 nm^2^). Thus, the actual accumulated exposure at the illuminated regions may be near 81 e^-^/Å^2^, while the average exposure across the entire crystal is likely an order of magnitude lower. This is evidenced by the high-resolution diffraction detected near the end of nanoEDT tilt series, which did not present an attenuation of diffracted signal commensurate with such a high electron exposure (Sup. Mov. 1 and 2). In fact, in conventional MicroED experiments^30^, significant radiation damage has been observed at electron exposures of as low as 3-10 e^-^/Å^2^.

We envision that integration of currently available hardware and software improvements, including improved angular sampling, cryogenic preservation procedures, precession of the probe and automation of crystal tilting, will greatly enhance the quality of data obtained by nanoEDT. Given the already high correlation of nanoEDT data to that collected by conventional MicroED methods, we see no absolute hinderance to the selective inclusion of diffraction from digitally defined regions of a sample. The similarity between nanoEDT and continuous rotation electron diffraction data (Fig. 4) indicates that nanoEDT may benefit from lattice variation due to nanocrystal bending. Lattice changes on the order of a few degrees have been detected across single nanocrystals^24^, whereby fixed-angle nanodiffraction averaged across large regions of a single crystal represent a pseudo-rocking curve more similar to a precession photograph than true fixed-angle diffraction.

Enabled by the control of nano-focused electron beams and sensitive detection of diffraction from nano-scale regions of single nanocrystals by direct electron detectors, nanoEDT has revealed the atomic structure of an amyloid-forming segment of the OsPYL/RCAR5 protein from digitally defined regions of single nanocrystals. Ultimately, the ability to selectively capture diffraction from digitally defined regions of a single nanocrystal or collection of nanocrystals (Fig. 5) could facilitate the unprecedented determination of atomic structures from heterogeneous or polycrystalline nanoassemblies.

## Materials and Methods

Detailed methods for OsPYL/RCAR5 peptide, nanoEDT data collection, nanoEDT data processing and phasing, MicroED data collection and processing and tomographic reconstruction are provided in the SI appendix.

## Author Contributions

M.G.J. and J.A.R. conceived of the project. J.A.R. directed the work. M.G.J. and S.L. grew, evaluated and optimized crystals. M.G.J., K.C.B., J.C., and J.A.R. collected data. M.G.J., C.O., and L.R. analyzed the data. M.G.J. and J.A.R. wrote the article, with input from all authors.

## Acknowledgments

We thank Drs. Duilio Cascio and Michael Sawaya (UCLA) for assistance with data processing and invaluable discussions, and Claudia Millan, Rafael J. Borjes and Isabel Uson (Institute of Molecular Biology of Barcelona) for facilitating use and implementation of the ARCIMBOLDO suite of programs. This work was performed as part of STROBE, an NSF Science and Technology Center through Grant DMR-1548924. This work is also supported by DOE Grant DE-FC02-02ER63421 and NIH-NIGMS Grant R35 GM128867. L.S.R. is supported by the USPHS National Research Service Award 5T32GM008496. S.L. is supported by the Next-Generation BioGreen 21 program (PJ01367602) through the Rural Development Administration, Republic of Korea. Work at the Molecular Foundry was supported by the Office of Science, Office of Basic Energy Sciences, of the U.S. Department of Energy under Contract No. DE-AC02-05CH11231. J.C. acknowledges additional support from the Presidential Early Career Award for Scientists and Engineers (PECASE) through the U.S. Department of Energy. J.A.R. is supported as a Searle Scholar a Pew Scholar and a Beckman Young Investigator.

