## Supplementary Information for "Atomic structures determined from digitally defined nanocrystalline regions"

**This PDF file includes:**

Supplementary Methods
Figures S1 to S7
Supplemental references

### Supplementary Methods

**Crystallization of AVAAGA.** Crystals of AVAAGA peptide, whose sequence was derived from residues 24-29 of OsPYL/RCAR5 (LOC_Os5g12260.1), were grown using the hanging drop method. OsPYL/RCAR5 is a positive regulator of ABA signal transduction in seed germination and early seedling growth from *Oryza sativa*. Lyophilised OsPYL/RCAR5 peptide (>98% purity by HPLC, GenScript) was dissolved in double distilled deionized water to a final concentration of 10 mg/ml. 2µl of the peptide was mixed with 2µl of a well solution consisting of 10% EtOH on a glass slide and equilibrated against 500 µl of well solution over a 24 well plate. High-quality, needle-shaped nanocrystals formed in 1 - 2 days.

**Sample preparation for diffraction experiments.** Three microliters of a crystal suspension were dispensed onto 400 mesh lacey carbon grids (Ted Pella) coated with either graphene oxide or 2 nm carbon films (UC type A) and allowed to adhere for 2 minutes before blotting excess solution and allowing to air dry.

**Collection of MicroED and fixed-angle selected area diffraction data.** Electron diffraction was carried out on a Tecnai F30 microscope operating at 300 kV. MicroED data was collected at liquid nitrogen temperatures while fixed angle diffraction was collected at room temperature. For microED data, a suitable crystal was identified in over-focused diffraction mode. The crystal was then isolated using a 1 µm selected area aperture and continuously rotated between 45 and -45 degrees at a rate of 0.3 degrees/second whilst being continuously illuminated by the electron beam. Diffraction frames were recorded as a movie using a TemCam-XF416 camera (TVIPS) with each frame corresponding to a 3-second exposure. For fixed-angle diffraction data collection, crystals were identified and isolated in a similar manner. Crystals were rotated between 45 and -45 degrees in discrete one-degree steps, and a 3-second exposure was recorded by the camera at each angular step. All measurements were performed at spot size 11 with the C2 lens set at 57% to ensure a low dose of ~0.01 $e^-/Å^2$/second or approximately 3 $e^-/Å^2$ per dataset.

**Collection of nanobeam electron diffraction tomography (nanoEDT) data.** Data collection for nanoEDT was performed on the TEAM I microscope at Lawrence Berkeley National Lab operating in microprobe STEM mode. The probe was focused to a size of approximately 12 nm with a semi-convergence angle of 0.09 mrad utilizing a 5 µm C2 aperture (Sup. Fig. 1). Samples were first located using a coarse STEM scan. Once a suitable crystal was located, the focused probe was raster scanned across the crystal with a step size of 40 nm covering a total area of approximately 1 by 3 µm. Data was recorded on a Gatan K2 direct electron detector operating at 400 frames per second, with each frame representing a single scan point. After each scan was completed the sample was rotated by 1 degree along the holder axis and the process repeated to give a final angular range of 45 to -45 degrees tilt. The total dataset then consisted of 30 by 70 by 90 individual diffraction patterns. Samples were maintained at liquid nitrogen temperature throughout data collection in a Gatan 636 holder. The total dose per frame was ~1 $e^-/Å^2$. The total accumulated dose is mitigated by having a step size significantly coarser than the probe size, thus reducing the likelihood that a specific sample volume will be illuminated with the same beam intensity at every tilt angle.

**Processing and data reduction of continuous rotation MicroED and fixed-angle selected area diffraction patterns.** MicroED data was converted to SMV format using the tvips-tools software package[1]. The fixed-angle selected area diffraction dataset was converted from tiff to binary files using a custom script in MatLab. All data were indexed and integrated using XDS[2] and merged using XSCALE.

**Processing and data reduction of nanobeam electron diffraction tomography data.** Raw data was first read into memory and preprocessed as previously described[3] (Gallagher-Jones et al., 2019) . In brief, raw data frames were aligned to a common center using the center of mass of the primary beam. The detector dark current was then subtracted from all frames using a median filter. A Gaussian model was fit to the distribution of pixel intensities after background subtraction to gain an estimate of the Gaussian noise of the detector. Using this model, a threshold was defined above which single or multiple electron counts were considered to have occurred. The values were separated into counting 'bins' using this threshold and the recorded values were converted to 'hybrid-counts'. The summation of these counts for all patterns within a single scan then represented the diffraction for that particular orientation.

To ensure that only diffraction frames deriving from the crystal were included in this sum, a virtual darkfield image was reconstructed at each scan. To do this a circular mask was defined and at each scan step all recorded electrons outside of this mask region (i.e. at high-resolution) were integrated in a manner analogous to recording with an ADF detector. In this darkfield image, pixels representing crystal regions were significantly brighter than those of the carbon support and so could be segmented via thresholding and morphological opening/closing (Sup. Fig. 1 & 2). The indices of the segmented pixels were then used to define which diffraction patterns in the scan would be combined or excluded. The summed diffraction patterns were then converted to binary files using a custom script in MatLab. Indexing and integration were performed in XDS and merged using XSCALE[2].

**Phasing and structure refinement.** The fixed-angle diffraction data and the MicroED dose series datasets, with the exception of the 12 $e^-/Å^2$ dataset, were phased by direct methods in SHELX[4] (Sheldrick, 2008) and refined using PHENIX[5]. Phases for the 12 $e^-/Å^2$ dataset were generated using the model from the 9 $e^-/Å^2$ in PHASER and then refined in PHENIX[5]. For the nanoEDT data, initial phasing was performed by a fragment-based search method using the ARCIMBOLDO software equipped with a library of poly-glycine 4mers derived from amyloid peptides, as described elsewhere (Richards *et al.* 2019). A 268-member library of tetrameric poly-glycine steric zipper fragments derived from over 100 previously determined structures were used in the program ARCIMBOLDO-BORGES[6,7]. Fragments were individually analyzed by Phaser rotation and translation analysis and top scoring chains were selected as inputs for SHELXE expansion by density modification and mainchain autotracing. The program was able to identify low mean phase error fragment solutions based on LLG and initial CC which were sufficient to provide initial phases despite failing to expand in SHELXE[4]. This fragment was then used as a starting point for building and refinement in COOT[8] and PHENIX[5] respectively.

**Estimation of crystal thickness from 4DSTEM data.** Crystal thickness was estimated using the log-ratio formula as employed in EELS experiments.

$$Z_{xy} = -\lambda \cdot \ln\left(\frac{I_{xy}}{I_0}\right)$$

Here $Z_{xy}$ represents the thickness at a given pixel and $\lambda$ is the mean free path of electrons through the peptide crystal, set at 332 nm as in previous experiments[3]. $I_{xy}$ is the transmitted intensity at a given scan position based on a combination of the integrated intensity of the central beam and the integrated intensity at Bragg peak locations identified from the aggregate diffraction pattern of the

4DSTEM scan. We found that omitting the electrons scattered elastically lead to erroneous over-estimation of transmission loss due to inelastic scattering. $I_0$ is estimated by taking the average value of transmitted intensity over vacuum (i.e. a hole in the lacey carbon) minus two times the standard deviation of the values in this region to account for fluctuations in the intensity of the electron beam.

**Tomographic reconstruction of crystals from virtual darkfield images.** Reconstructed maps of crystal thickness as described above were first roughly aligned to a common tilt axis using features of the lacey carbon substrate. Due to sample drift during data collection, the images were cropped to remove any regions of the crystal that were not consistently in the field of view throughout the entire tilt series. A flat background was calculated from regions of the images that sampled vacuum and subtracted and any resulting negative values were set to zero. Tomographic reconstruction was performed using the GENFIRE algorithm with image shift refinement[9]. Reconstructed volumes were visualized with CHIMERA[10].

**Comparison of integrated intensities.** Intensities were merged together using Scalepak[11] to ensure that only common reflections were compared. Fourier magnitude plots and linear regression were calculated in MatLab and comparisons of zone-axis reflections was performed using viewHKL[12].

**Peak identification in multi-lattice data.** To enhance the contrast of the multi-lattice diffraction patterns the data was binned by 5. Peaks were then localized via template matching with a circular template 6 pixels in diameter using normalized cross correlation implemented in MatLab.

**Code Availability**

The MatLab scripts for data pre-processing can be found at: https://github.com/marcusgj13/4DSTEM_dataAnalysis. The MatLab implementation of GENFIRE used for tomographic reconstructions can be found at:
https://github.com/apryor6/GENFIRE-refine3D

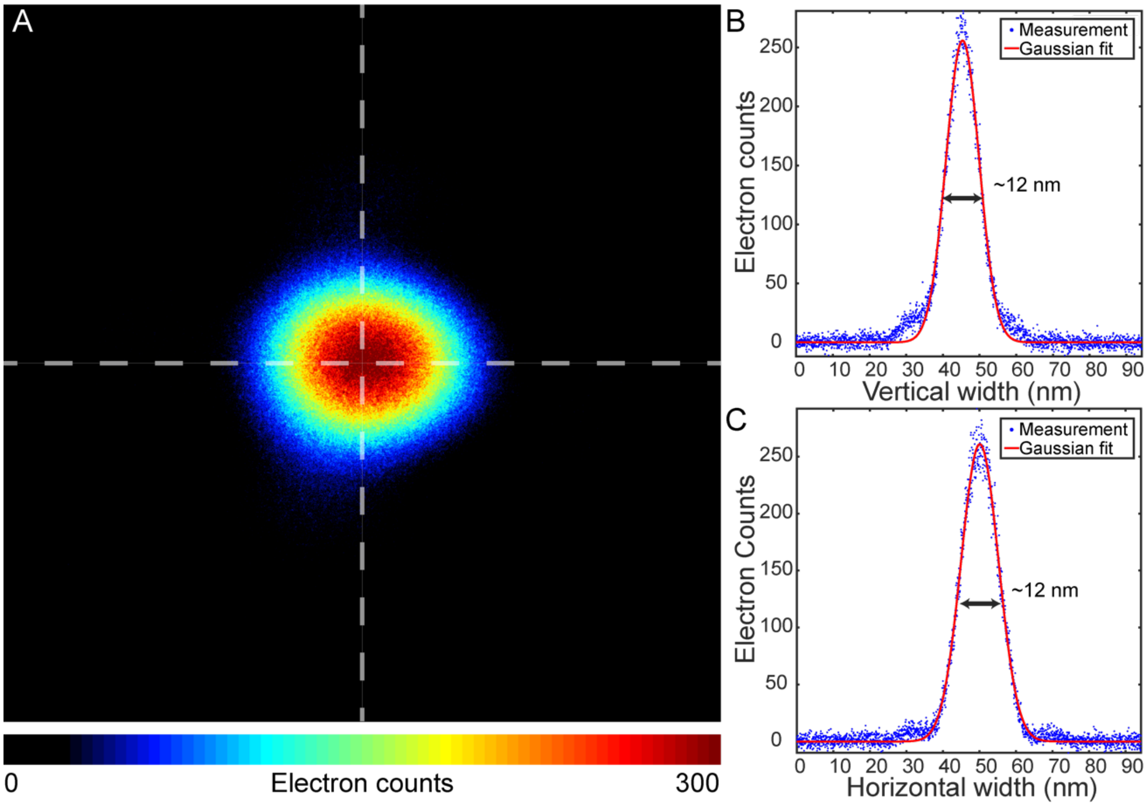


**Sup. Fig. 1.** Analysis of the electron probe used in nanoEDT experiments. (A) Image of the focused probe captured on a Gatan Ultra scan 1000 with an exposure time of 3s. (B) Vertical cross-section of the probe (vertical white line in A) and Gaussian fit to the profile giving an estimated diameter of ~12 nm at FWHM. (C) Horizontal cross-section of the probe (vertical white line in A) and Gaussian fit to the profile giving an estimated diameter of ~12 nm at FWHM.

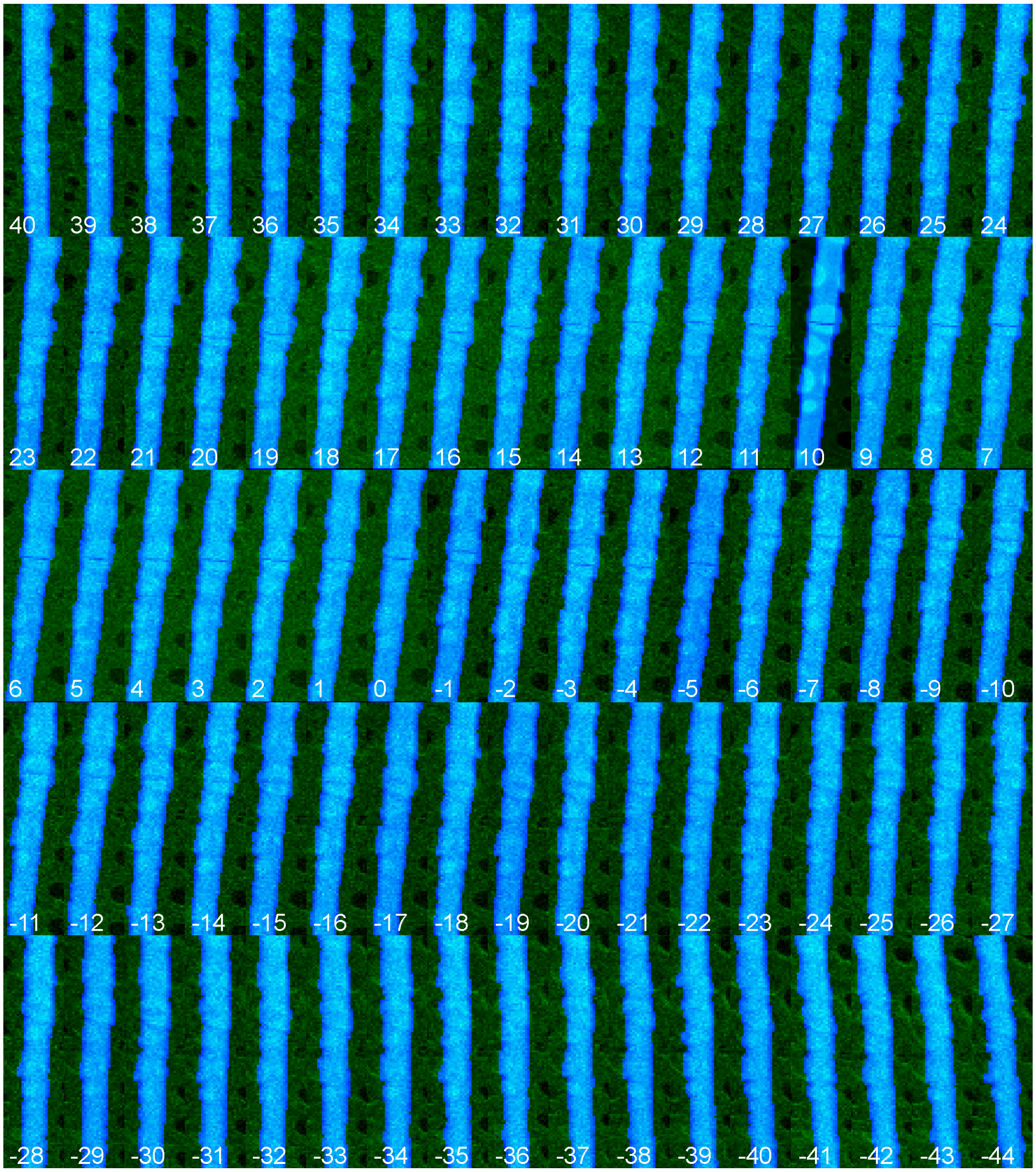


**Sup. Fig. 2.** Digitally defined regions of nanoEDT scans taken from the first crystal. Blue pixels represent scan locations that were used to assemble diffraction patterns for the first dataset. Green and black pixels were excluded.

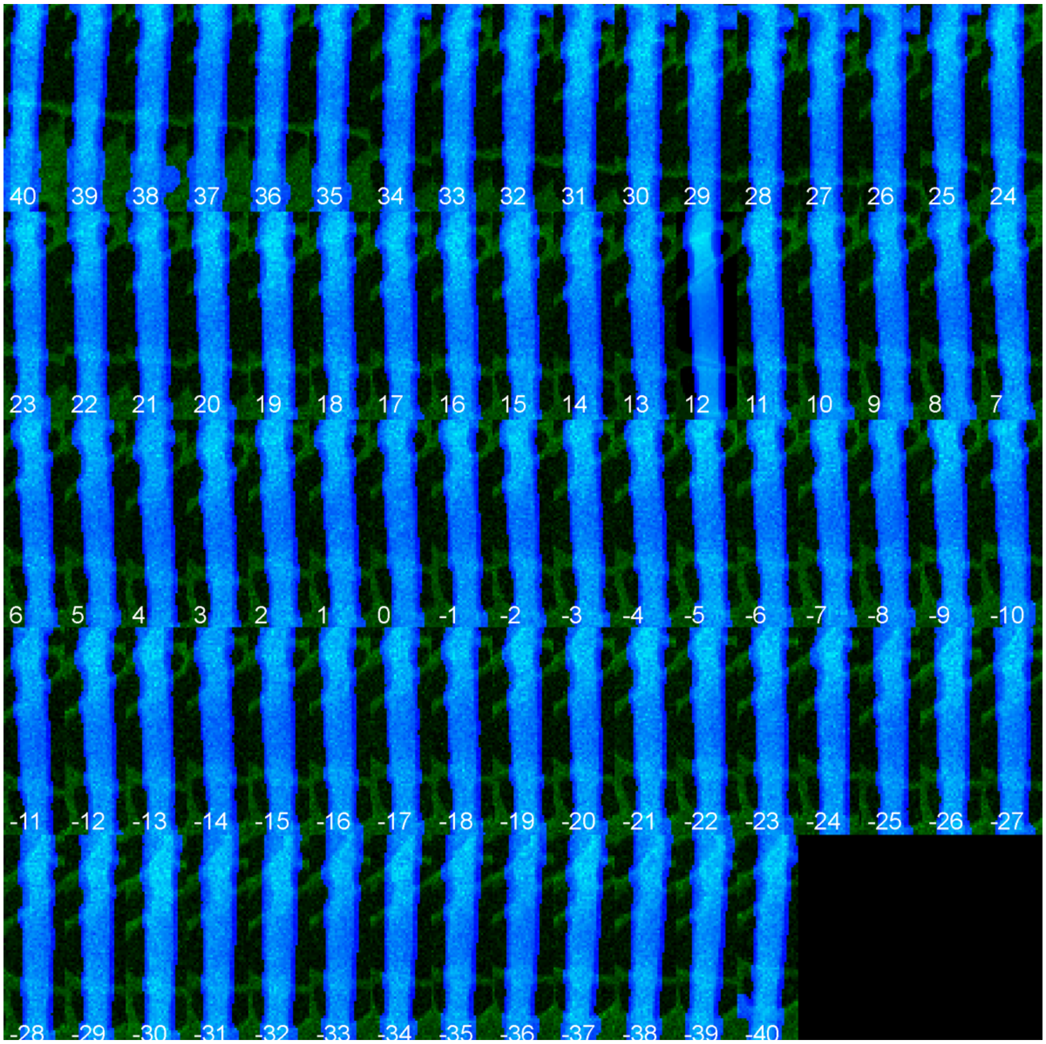


**Sup. Fig. 3.** Digitally defined regions of nanoEDT scans taken from the second crystal. Blue pixels represent scan locations that were used to assemble diffraction patterns for the second dataset. Green and black pixels were excluded.

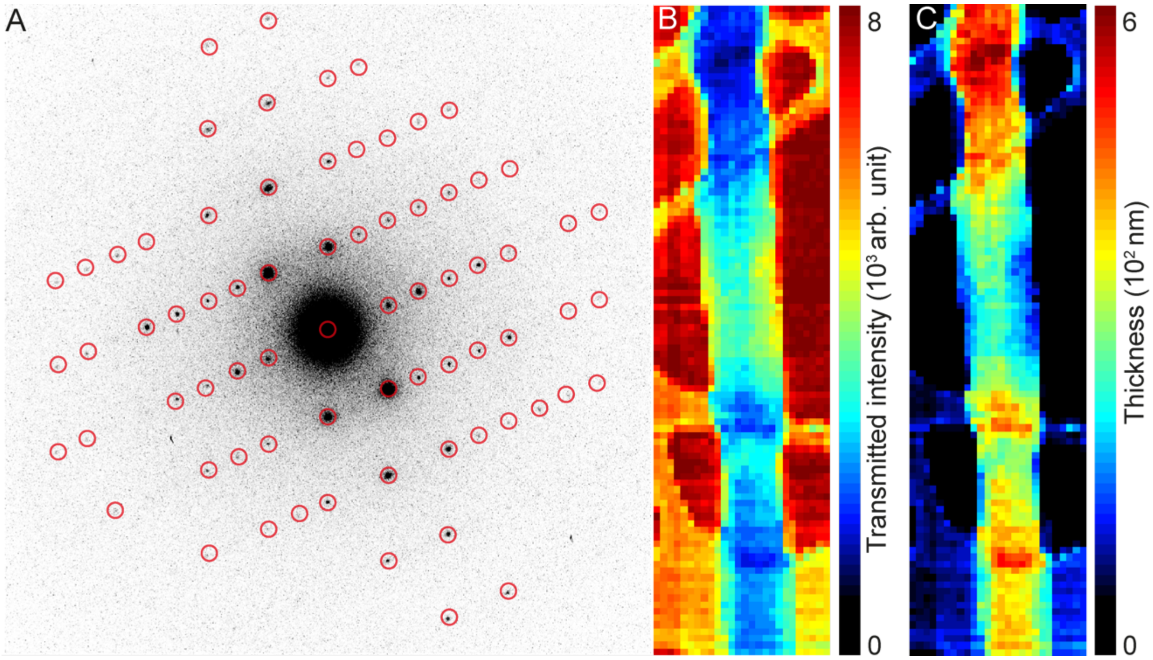


**Sup. Fig. 4.** Estimation of crystal thickness from 4DSTEM data. (A) Aggregate diffraction pattern calculated from a 4DSTEM scan. Highlighted by red circles are the regions of the diffraction pattern used to estimate changes in transmission of the central beam due to inelastic scattering. (B) Map of transmission loss due to inelastic scattering across the 4DSTEM scan calculated from highlighted regions in (A). (C) Estimated thickness of the crystal and surrounding regions based on changes in transmission in (B).

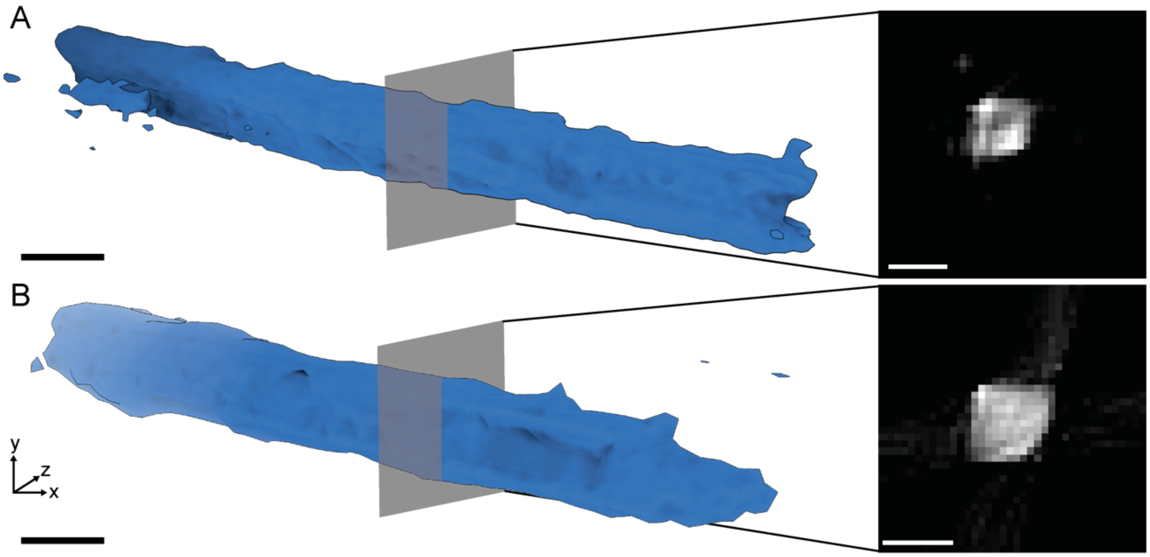


**Sup. Fig. 5.** Tomographic reconstruction of crystals from calculated thickness maps. (A) Isosurface rendering of the tomographic reconstruction of the first crystal analyzed by nanoEDT in this study. The gray plane represents a slice through the reconstructed density. (B) Isosurface rendering of the tomographic reconstruction of the second crystal analyzed by nanoEDT in this study. The gray plane represents a slice through the reconstructed density. All scale bars represent 400 nm.

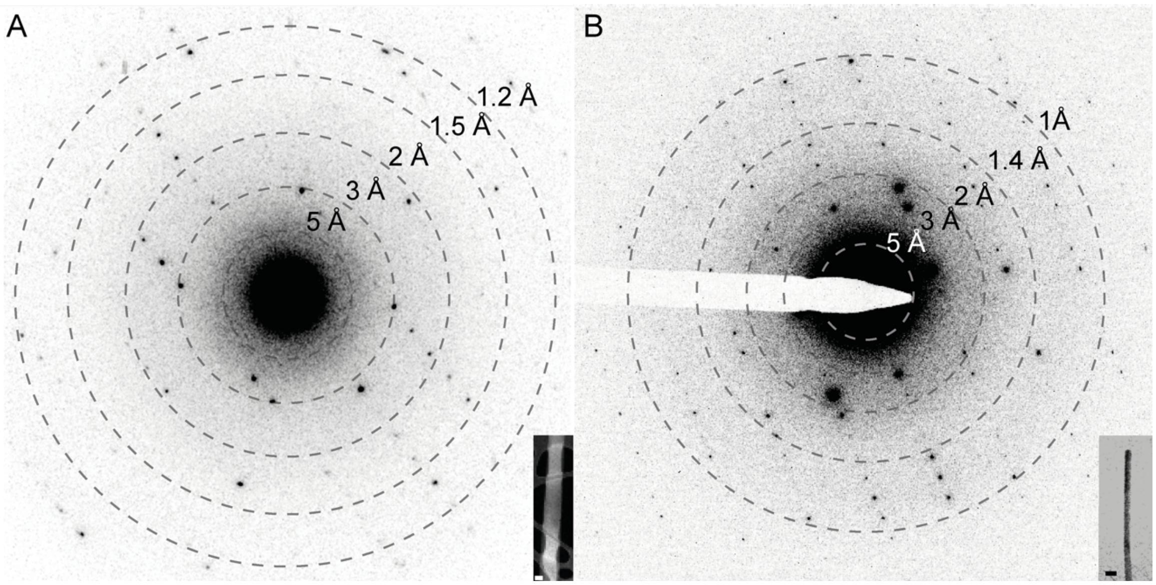


**Sup. Fig. 6.** Diffraction patterns obtained by nanoEDT *vs.* conventional MicroED. (A) Computed nanoEDT diffraction pattern from 4DSTEM scans recorded on a K2-IS detector. (B) MicroED diffraction pattern recorded on a TemCam-XF416 CMOS detector. Inset are the crystals that the diffraction was captured from. Scalebars are 200 nm.

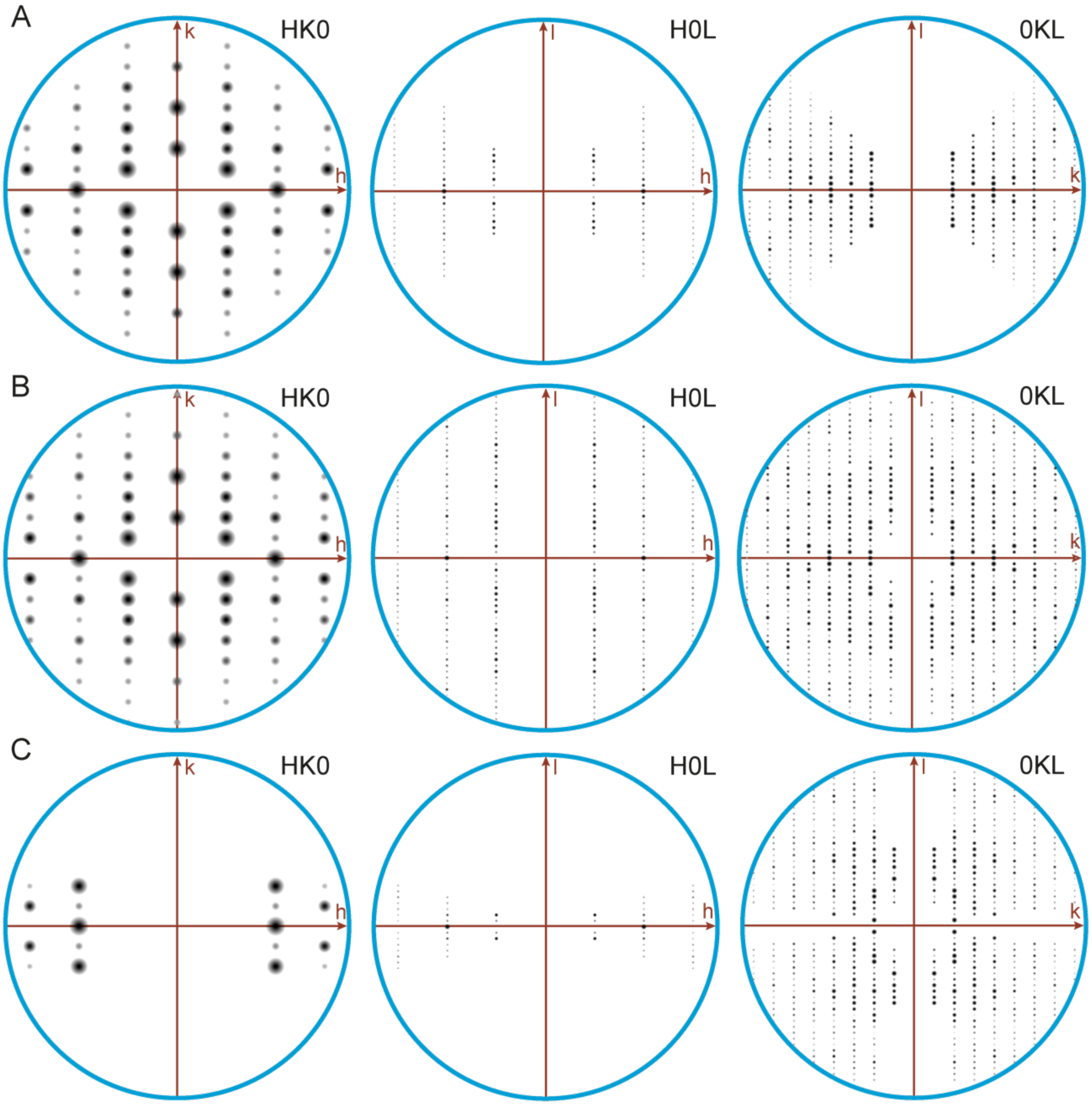


**Sup. Fig. 7.** Comparison of Integrated intensities at principle zone axes. (A) nanoEDT. (B) MicroED dataset. (C) fixed-angle SAED dataset
